# The causal effect of adiposity on hospital costs: Mendelian Randomization analysis of over 300,000 individuals from the UK Biobank

**DOI:** 10.1101/589820

**Authors:** Padraig Dixon, William Hollingworth, Sean Harrison, Neil M Davies, George Davey Smith

## Abstract

Estimates of the marginal effect of measures of adiposity such as body mass index (BMI) on healthcare costs are important for the formulation and evaluation of policies targeting adverse weight profiles. Many existing estimates of this association are affected by endogeneity bias caused by simultaneity, measurement error and omitted variables. The contribution of this study is to avoid this bias by using a novel identification strategy – random germline genetic variation in an instrumental variable analysis – to identify the presence and magnitude of the causal effect of BMI on inpatient hospital costs. We also use data on genetic variants to undertake much richer testing of the sensitivity of results to potential violations of the instrumental variable assumptions than is possible with existing approaches. Using data on over 300,000 individuals, we found effect sizes for the marginal unit of BMI more than 50% larger than multivariable effect sizes. These effects attenuated under sensitivity analyses, but remained larger than multivariable estimates for all but one estimator. There was little evidence for non-linear effects of BMI on hospital costs. Within-family estimates, intended to address dynastic biases, were null but suffered from low power. This paper is the first to use genetic variants in a Mendelian Randomization framework to estimate the causal effect of BMI (or any other disease/trait) on healthcare costs. This type of analysis can be used to inform the cost-effectiveness of interventions and policies targeting the prevention and treatment of overweight and obesity, and for setting research priorities.

## 1 Introduction

A positive association between adiposity and healthcare costs is well established. It has been documented for a variety of different contexts, circumstances and health systems (1–3). This association has powerful economic salience because of its apparent consequences for the level, growth and composition of healthcare spending.

The underlying biological relationship between adiposity and health is complex (4), but the endocrinal (5), cardiometabolic (6, 7) and other changes (8) associated with increased adiposity are themselves linked to substantial healthcare resource requirements (9). Increases in the mean and variance of adiposity, reflected in widely used measures of nutritional status such as body mass index (BMI - weight divided by the square of standing height) have led to important changes in the global distribution of adiposity (10–12). The worldwide prevalence of overweight (BMI>=25kg/m^2^) and obesity (BMI>=30kg/m^2^) is 28.8% for men and 29.8% for women. This accounts for some 2.1 billion individuals, an increase of approximately 50% since 1980 (13). More individuals globally are now either overweight or obese than are underweight (11, 14).

Correlational evidence of the BMI-cost association is influential. Examples of this influence include the development of guidelines and policies to prevent obesity (15), evaluation of interventions targeting overweight and obesity (16), and the prioritization of research into the consequences of obesity (17). However, a critical limitation of much if not all of this multivariable^1^ research is that it can be seriously affected by endogeneity bias (18).

This endogeneity arises through three channels. The first is measurement error arising from mismeasurement of BMI (and other measures of adiposity), particularly where individuals selfreport weight (19, 20). The second is reverse causation or simultaneity bias, which would occur if healthcare costs influenced adiposity. The third source of bias is omitted variable bias, arising from unknown or unmeasured common causes of both adiposity and healthcare costs.

The direction of the omitted variable bias will generally not be known a priori. Disease processes that are related to healthcare costs may also influence adiposity. For example, higher BMI is associated with increased risk for cancers (21), but cancer (including prodromal cancer) may itself lead to reductions in BMI (22). Similarly, people with higher BMI are more likely to smoke, while smoking itself lowers BMI (23, 24). Without evidence of the wider determinants of both adiposity and healthcare cost, the analyst cannot reliably predict directions of bias when undertaking multivariable analyses of this association.

BMI-health outcome associations are therefore distorted because one of the drivers of health outcomes like health care cost is own health status. This observation has motivated attempts to use instrumental variable (IV) analyses in which the instrument for own BMI is the BMI of a biological relative, for example in relation to the association between BMI and mortality (25). This approach has also been used to model the causal impact of adiposity on costs, and arguably represents the most credible attempt to date to overcome the endogeneity biases of multivariable analysis.

For example, Cawley and Meyerhoefer (26) used the BMI of a biological relative as an IV. This suggested that the healthcare costs of obesity were drastically underestimated by prior multivariable analyses, with a fourfold difference in the marginal costs of obesity between multivariable and causal IV analysis reported, and a threefold difference in the costs of a marginal unit of BMI. Large but less pronounced differences between multivariable and IV models were also reported in studies using similar instruments by Cawley et al (27), Black et al (28) and Kinge and Morris (29).

However, this approach does have limitations. The association of biological relatives and healthcare costs may itself be affected by omitted variables that are common and independent causes of both BMI and healthcare costs. These could include the home environment that is shared by biological relatives and which may influence food consumption, proclivity to exercise, and access to and use of healthcare services. People who have children (necessary for the biological relative approach) may differ from those who do not have children. Intrauterine influences of maternal BMI on offspring BMI, such as smoking and alcohol drinking during pregnancy (30), and genetic influences that affect healthcare costs other than through adiposity (31), will also confound this relationship.

This paper exploits a novel identifying approach - germline genetic variation associated with BMI – in an instrumental variable analysis. This approach has the advantage (in principle) of avoiding the limitations of both multivariable analysis and the use of a biological relative as an instrument. At each point of variation in the genome, offspring typically inherit one allele from their mother, and one from their father. This random inheritance of alleles is a natural experiment, in which individuals in a population can be divided into groups based on their inherited “dosage” of these variants (32). If the instrumental variable assumptions hold, these genetic variants can be used to test whether BMI affects healthcare costs. Using genetic variants as IVs in this way has become known as Mendelian Randomization (33).

Robust evidence of the causal association between adiposity and healthcare costs is a critical input for the formulation and evaluation of cost-effective policies and interventions targeting (in particular) overweight and obesity (8), as well as for identifying research priorities in this area. The widespread use of models lacking robust identification may substantially underestimate the true causal effects of obesity. Very large, high-quality datasets that can facilitate this type of analysis are beginning to become available (34, 35) but remain largely if not entirely unexploited by health economists studying the causal effect of health conditions and traits on cost outcomes. Below, we introduce Mendelian Randomization as a form of IV analysis.

## 2 Methods

### 2.1 Mendelian Randomization and instrumental variable analysis

Here, we briefly introduce the high-level biological mechanisms that motivate the use of genetic variants in IV analysis. More detailed introductions and extended overviews of Mendelian Randomization are available elsewhere (36–39).

A single nucleotide polymorphism (SNP) is a specific location (or locus) in the human genome that differs between people in the population. At each SNP people will have two alleles, one for each chromosome. During cell division at conception (meiosis), offspring inherit at random one of their mother’s two alleles, and one of their father’s two alleles. Specific SNPs or sets of SNPs are known to associate with particular health conditions or influence the development of particular traits. Thus, the phenotype (a measurable disease or trait such as BMI) may be influenced by genotype (an underlying genetic structure associated with the phenotype).

The provenance of the term Mendelian Randomization (40), and the potential utility of genetic variants as IVs, is founded on Mendel’s first and second laws of inheritance. The first law describes random segregation of alleles from parent to child during the formation of gametes. The second law describes the independent assortment of alleles for different phenotypes at conception. Genetic variants that are in different locations in the genome are generally inherited in a way that is independent of the inheritance of other genetic variants. The allocation of these genetic variants to offspring is therefore random, conditional on parental genotype.

We now describe the core instrumental variable assumptions in the context of Mendelian Randomization. These assumptions can be described as comprising the relevance assumption, the independence assumption, and the exclusion restriction.

The first IV assumption (“relevance”) is that the instrument should be associated with the treatment variable, which in the case of this paper is BMI.^2^ The associations of SNPs with diseases and traits are readily determined from genome wide association studies (41, 42), which study the independent association with specific phenotypes of many SNPs - potentially hundreds of thousands - across the genome. These associations are corrected for multiple testing so that genome-wide significance is obtained as the conventional p-value threshold value based on an alpha of 0.05 divided by *k*, where *k* can be interpreted (conservatively) as the number of independent statistical tests conducted across the genome (43) if there are one million independent genetic variants (44). These associations will be validated in independent replication samples (45). Following convention, we will describe p <= 5×10^-8^ as genome-wide significant.

The second assumption is that there are no omitted variables in the associations of the IV and the outcome (health care costs). This assumption is generally unproblematic since the SNPs are determined at conception, and therefore prior to the postnatal circumstances, events and behaviours of later life. However, time of conception (such as month or year of birth) could theoretically associate with SNPs and health care costs. Population stratification, the separation of individuals into distinct subgroups that differ in allele frequencies, is one means by which the second assumption may be violated, since differences in alleles in this case would indicate differential ancestry rather than disease susceptibility (46).

Ancestry influences the distribution of genetic variants, but also risks of disease not necessarily due to those variants. This potential confounding by ancestry is typically accounted for by adjusting for the genetic principal components (47) and restricting analysis to genetically homogenous ethnic groups. Simultaneity bias, if present at all and absent population stratification, is likely to be modest although this assumption is best tested rather than assumed for any particular study (48). Examples of independence of common genetic variation from common omitted variables (and thus that SNPs are likely to be independent of environmental influences) has been demonstrated empirically (37, 49).

The third IV assumption is that the SNP(s) affect the outcome only via the treatment variable; that is, via the condition or trait of interest. This is the exclusion restriction. Violations of this assumption is the primary threat to the validity of IVs used in Mendelian Randomization. There are two principal mechanisms by which this assumption may be violated in Mendelian Randomization.

The first is the correlation of the SNP(s) in question with other SNPs that affect the outcome through a path other than via the condition or trait of interest (50). This correlation of variants, known as linkage disequilibrium, arises when particular variants tend to be inherited together (contrary to Mendel’s second law), generally because they are located in close physical proximity on the genome (51).

The second mechanism concerns variants that affect more than one phenotype. A SNP that affects BMI may also, for example, affect the risk of depression through a BMI–independent mechanism. IV analysis relating, for example, a set of BMI SNPs to cost outcomes would suffer from omitted variable bias in this case if depression independently affects both BMI and healthcare costs. This is sometimes known as horizontal pleiotropy (37). Pleiotropy (52, 53), the effect of a single SNP on multiple phenotypes (53, 54), may be pervasive throughout the human genome (55). There would be no bias in this analysis if depression was on the causal pathway between BMI and healthcare costs, a situation sometimes referred to as vertical pleiotropy (37) (37), or if the other phenotype did not affect the outcome of interest.

Finally, we note that Mendelian Randomization does not identify population average treatment effects. IV analysis identifies local average treatment effects if a monotonicity condition holds. Monotonicity requires that the direction of effect on the treatment from varying the level of the instrumental variable should be in the same direction for all individuals. When monotonicity is satisfied, IV analysis (including Mendelian Randomization) identifies a local average treatment effect; that is, an effect in those whose treatment would differ if the value of the IV differed.

### 2.2 Estimation

For a single SNP, the ratio (or Wald) estimator can be calculated as the ratio of the association of the outcome with the variant to the association of the treatment variable (BMI in this paper) with the SNPs. This gives the effect of the variant on the outcome, scaled by the effect to the SNPs on the treatment. This is equivalent to the two-stage least squares estimator for a single SNP. Using the terminology of Bowden et al (56), indexing individuals by *i* and denoting SNPs as G (indexed *j* from 1 up to *J*) these two relationships can be written as:

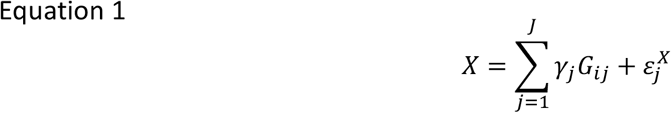

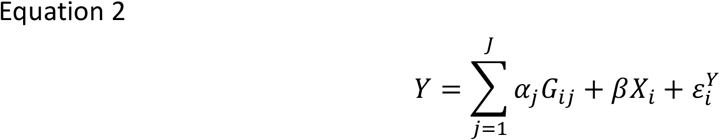

Without loss of generality, we ignore constants and exogenous omitted variables in Equations 1 and 2. The alpha term is the direct effect of variants on the outcome that do not operate through the BMI treatment variable. If the exclusion restriction holds then alpha will be zero, since valid instruments influence outcomes only through an effect on the treatment.

Note also that the two associations described in Equations 1 and 2 in the Wald estimator need not come from the same sample, in which case a two-sample IV estimator is used (57). A two-sample approach using summarized data may offer similar or better efficiency than a single sample study using individual-level data (58), particularly if larger sample sizes are available under a two-sample approach. In the two-sample setting, genetic variants should have similar effects in each population. One simple practical test for this in Mendelian Randomization is to examine similarity in terms of ethnic group and distributions of sex and age (48).

Rewriting equations (1) and (2) into the reduced form yields:

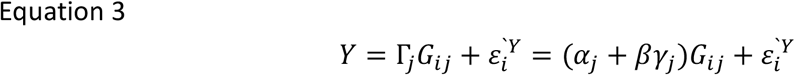

Where 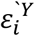 is the error term of the reduced form. The ratio estimate is ratio of the effect of the SNPs on the outcome, scaled by their effect on the treatment, which can be written (ignoring the error term) as:

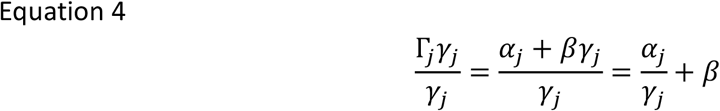

The ratio estimates from each individual variant can be combined using weighted regression or equivalently inverse variance weighted (IVW) meta-analysis to produce an overall causal estimate (henceforth for simplicity we refer to this estimate as the IVW estimate – Equation 5).

This assumes that there no correlation between the Wald estimates for each SNP, which will hold if they are not in linkage disequilibrium.
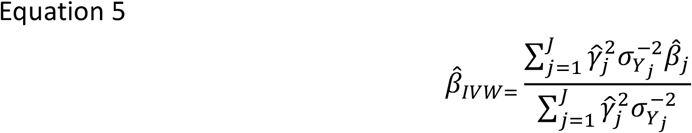

Here, the 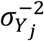 terms are the variance of the error term in the outcome-SNP regression models; the small variance of the error term in the treatment-SNP regression is ignored (the no-measurement error assumption).

The estimates for each variant should converge toward the same causal parameter estimate if the exclusion restriction holds; put differently, there should be no more heterogeneity in the estimates for all parameters than would be expected by chance. This can be assessed using Cochran’s Q statistic (59, 60) in two-sample settings (this is closely related to Sargan’s over-identification test (61) in single sample settings), which follows a χ^2^ distribution with *J*-1 degrees of freedom:

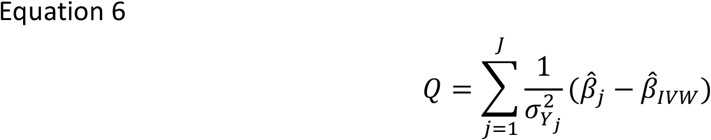

Cochran’s Q can identify failure of the instrumental variable assumptions, but not whether this is due to one, some or all IVs being invalid. As such, it is a relatively crude “catch all” test of instrument validity.

### 2.3 Sensitivity analysis

A number of methods have been developed to accommodate violations of the exclusion restriction due to pleiotropy that is suggested by (but not necessarily unambiguously identified by) high heterogeneity as indicated by Cochran’s Q (62, 63). The following considers different methods for assessing possible violations of the exclusion restriction, in the spirit of Conley et al (64), in relaxing the assumption that the alpha parameter is exactly zero and in seeking methods to generate consistent estimates of the causal effect even if some or all of the IVs are invalid.

If pleiotropy (i.e. non-zero *α_j_* terms) is present but small in magnitude then biases in any causal analysis will be modest. If *α_j_* is zero on average across all SNPs then the relationship is estimated with more noise and hence some loss of efficiency than if all *α_j_* values were zero, but the bias term will have zero mean on average even if some or all of the pleiotropic effects are large. In this case, the IVW estimator could be implemented using a random effects metaanalysis.

If the mean effect of alpha is not zero, then directional pleiotropy is present. So-called MR-Egger methods allow for directional pleiotropy by modelling both the slope and intercept of the ratio estimator.
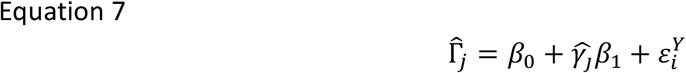

Note that the “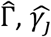” terms included in Equation 7 are themselves estimates, respectively the SNP-cost and SNP-BMI estimates. All SNPs can be invalid instruments under MR-Egger, provided that the InSIDE assumption (<u>In</u>strument <u>S</u>trength <u>I</u>ndependent of <u>D</u>irect <u>E</u>ffect) assumption holds. The MR-Egger effect estimate can be written as 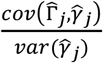 which can be re-expressed as the true effect estimate 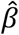 plus a bias term 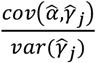. This InSIDE assumption appears to be plausible in some cases (e.g.(65)) but less so in others (e.g (37, 56)). This assumption is most likely to hold in this context when it is the exclusion restriction assumption that is violated due to horizontal pleiotropy, rather than to other violations of the IV assumptions, such as associations with omitted variables in the BMI-cost relationship. MR-Egger estimators are less powerful and less efficient than the estimators discussed below because of the need to estimate both the slope parameter and the intercept parameter.

An alternative to relying on the InSIDE assumption is to use the median ratio estimate of all available instruments (66). This estimator will be unbiased if more than half of the instruments are valid, i.e. *α_j_* = 0 for at least half of all SNPs. The simple intuition for this estimator is that invalid instruments in the IVW approach will contribute weight to the overall regression estimate and will be biased even asymptotically. On the assumption that the majority of instruments are valid, then invalid instruments contribute no weight and are less biased than IVW in finite samples and unbiased asymptotically. We implement a penalized weighted median estimator. SNPs contributing to the median 50% of the statistical weight are used to form the median estimate. The weights are a function of the precision with which SNPs are estimated, and the penalization involves “down weighting” outlying SNPs that contribute substantial heterogeneity to the Cochran Q statistic.

The final class of estimators we consider are mode based (67). The underlying assumption, in terms of the alpha expression, is that *Mode*(*α*_1_, *α*_2_, … *α_k_*) = 0. The intuition is that classifying variants into clusters based on similarity of effect will be consistent if the largest homogenous cluster are valid SNPs. All other SNPs outside this cluster, even a majority of individual variants in the sample, could be invalid, provided this “zero model pleiotropy” assumption holds. This approach requires the setting of an arbitrary bandwidth parameter to define the clusters. We implement a more efficient version of the simple median estimator by weighting median estimates by the inverse variance of the effect of the SNPs on the outcome. This is given effect by creating an empirical density function formed from the weighted mode estimates.

It is important to note that the second and third IV assumptions are not directly testable, and the assumptions underlying alternative modelling approaches for *α_j_* term are themselves untestable. However, these approaches are important forms of sensitivity analysis that allow the instrumental variable assumptions to be relaxed, albeit at the cost of other untestable assumptions. Similarity of estimated effect under each of the estimators considered would offer some reassurance that the same causal effect is being identified, although MR-Egger offers much lower precision than the other estimators.

We also assessed whether any heterogeneity present in the main analysis was also present when analyzing broad categorization of overall inpatient hospital costs into elective costs, nonelective costs, and other costs. Elective admissions are those that are planned. Non-elective admissions are not arranged in advance, and may include, for example, emergency care or maternity care. The “other” category includes all other costs, including day cases, which are elective admissions but where an overnight stay is neither planned nor undertaken. The inverse-variance weighted Mendelian Randomization models were re-run using only costs in each respective category as the outcome variable. We examined effect sizes, p-values and heterogeneity statistics for each model and compared them to the base “all inpatient costs” analysis. We emphasize that the distinctions between these sub-categories are not absolute, since the categorization is somewhat arbitrary. More details are provided in the Supplementary Material.

We performed four further sensitivity analyses. The first considered whether the association between BMI and healthcare cost may be non-linear, the second estimated a multivariable Mendelian Randomization instrumenting for both BMI and body fat percentage, the third assessed the impact of potential weak instrument bias, and the fourth involved a within-family Mendelian Randomization analysis to avoid bias from so-called dynastic effects (68).

#### Non-linear models

There is some evidence of a non-linear association between BMI and hospital costs from multivariable and causal studies (e.g. (26, 27)), although the presence or absence of such an effect can reflect the specific modelling approach used. Fitting non-linear models in the IV settings is complicated when the instruments explain a relatively small proportion of variance in the treatment (as in the present example), because any non-linear effects may not be detectable in the relatively narrow range over which such effects influence the treatment-outcome association (69).

To avoid this, we used methods developed by Staley and Burgess (69) to test for and model non-linearity in Mendelian Randomization models. This approach involves the following steps. First, a control function approach is adopted in which the IV-free distribution of BMI is estimated. This is necessary because stratifying on the original distribution may induce an association between the IV and the cost outcome that violates the exclusion restriction – this possibility arises because BMI is itself a potential outcome intermediate between the IV and the cost outcome. The second step involves choosing the number of quantiles into which BMI is to be stratified, before the estimation of the local average treatment effect in each stratum. The estimation of the treatment-outcome model from these data can then be implemented using either a fractional polynomial method and a piecewise linear method. We implemented the former method, based on an assumption that the relationship between BMI and costs is more likely to be relatively smooth than piecewise in the type of sample analyzed.

#### Multivariable Mendelian Randomization – BMI and body fat

Multivariable Mendelian Randomization can estimate the direct causal effect of more than one treatment (70, 71) and therefore allows potentially pleiotropic exposures to be jointly modelled to avoid violations of the exclusion restriction. For this analysis we remain agnostic as to which of the two measures of adiposity that we study below- BMI and percentage of body fat – more accurately index the health-compromising consequences of fatness. The percentage of body fat arguably better captures body composition than does BMI and may better predict particular health outcomes (e.g.(20, 72, 73)), but BMI nevertheless retains broad applicability and utility as an easily measured variable that offers robust associations with a variety of relevant health outcomes (4).

In this application of multivariable Mendelian Randomization, genetic variants for BMI and for percentage of body fat are included in the same model. This allows for these biologically related treatments to be modelled together, and for the potential mediation of one treatment (BMI for example) by another (body fat percentage) to affect the outcome. The coefficients in the estimated models reflect the direct causal effect of each treatment, holding the other treatment fixed. These models have considerably lower power to detect causal effects than univariable Mendelian Randomization, but the analysis can nevertheless usefully estimate the direct effect of BMI on outcome compared to the total (comprising the direct effect of BMI and its indirect effects via body fat percentage) estimated in conventional Mendelian Randomization (74).

#### Weak instruments

We estimated the “robust adjusted profile score” model of Zhao et al (75), which is unbiased in the presence of many weak instruments, and is also robust to measurement error in the SNPs-treatment models. Even if SNPs satisfy the relevance assumption at genome-wide levels of significance, it is possible that they are “weak” instruments, in the sense of explaining only a small proportion of the variance in the treatment (76, 77). Weak instruments will bias the causal estimate in finite samples toward the non-IV estimate (76, 78). The Zhao et al approach relies on a version of the InSIDE assumption that underpins the MR-Egger approach, but unlike MR-Egger assumes that the pleiotropic effects *α*_1_, *α*_2_, … *α_k_* have mean zero.

#### Within-family Mendelian Randomization

Finally, we consider within-family Mendelian Randomization. This is intended to address biases from dynastic effects (68), although within-family analysis also avoids any biases caused by cryptic population structure not accounted for by restricting analysis to homogenous ethnic group and the use of genetic principal components. Dynastic effects refer (in the present context) to the direct effect of parents’ BMI on their children. This type of effect may reflect non-transmitted alleles (79) – even if children do not receive a BMI-increasing SNP, their parents may possess such a SNP and this in turn can influence the environment in their children are raised. If present, the main Mendelian Randomization analysis presented here would wrongly attribute some of the influence of parental BMI to the child’s BMI-increasing SNPs that are included in the analysis. We therefore explored whether bias from dynastic effects could be reduced by conducting a within-family Mendelian Randomization in which a family “fixed effect” adjusts for environmental conditions created by parents that are shared by offspring (38).

Siblings were identified in the UK Biobank by using data on kinship taken from the KING toolset and data on the proportion of loci shared between individuals. More details are available in Brumpton et al (80). We restricted analysis to the IVW estimator. This is because the MR-Egger, median and mode estimators used in the main analysis on the sample of unrelated individuals have lower power than the IVW estimator. The sample of included related individuals is 11% (n = 36,196) of that used in the main analysis, and the power of IVW methods is therefore much reduced. We estimated fixed effect instrumental variable models, clustering on family units, and conditioning on sex and the first ten genetic principal components.

## 3 Data

### 3.1 UK Biobank

Individual-level data were drawn from the UK Biobank study. This large prospective cohort enrolled 503,317 adults (representing a response rate of 5.45%) aged between 37 and 73 (99.5% of enrollees were aged between 40 and 69) living in England, Scotland and Wales (81). At the baseline appointment, participants completed a number of questionnaires, biomarker specimens were drawn, physical function was assessed, and consent was given to link these data to death registers and healthcare records (34).

Weight and height were measured at the baseline appointment by nurses. Weight was measured using weighing devices. Body composition was measured using bio-impedance (opposition of alternating current to adipose tissue). Both measures were very similar (Lin’s rho p-value < 0.001) and impedance-based BMI data were used when the conventional BMI data was missing. Observations that had a mean difference between traditional and impedance-based measures of BMI of more than 5 standard deviations from the mean difference were excluded from the analysis. Whole body percentage fat mass calculated from impedance was used in the multivariable Mendelian Randomization analysis.

### 3.2 Measurement of costs

Admitted patient care episodes, sometimes referred to as inpatient care episodes, were obtained from Hospital Episode Statistics (HES) (for English care providers) and from the Patient Episode Database for Wales (for Welsh providers) that were linked to UK Biobank. Inpatients are those admitted to hospital and who occupy a hospital bed but need not necessarily stay overnight (i.e. day case care). Due to differences in the collection and valuation of care in Scottish hospitals compared to hospitals in England and Wales (82), only costs from the latter two jurisdictions are included in this analysis. Linkages to other forms of care were not available at the time of writing.

Each “Finished Consultant Episode” (FCE) can be characterized by a number of variables, most importantly procedure codes (83) and diagnosis codes (based on ICD-10 codes (84)). These FCEs were converted, using NHS software (85), into Healthcare Resource Groups (HRGs). HRGs are used for casemix-adjusted remuneration of publicly-funded hospitals in England and Wales. Unit costs were assigned to each HRG, and inpatient costs per person year of follow-up were calculated for each patient on the basis of their recorded FCEs (if any). Further details on the cost calculations are given in Dixon et al (86).

Only episodes and UK Biobank baseline appointments occurring on or after 1 April 2006 were eligible to be included in the analysis because of changes to the hospital payment system that came into effect at that time (87). Data on inpatient episodes was available until patient death, patient emigration (rates of which are estimated to be a modest 0.3% (81)), or the censoring date for inpatient care data of 31 March 2015. Cost data are reported in 2016/17 pounds sterling.

### 3.3 Genetic data and linkage to phenotypic data

Genetic data was subject to quality controls by UK Biobank (88), as well as further in-house processing and management (89, 90). Briefly, 488,377 individuals in the UK Biobank were successfully genotyped. Removal of individuals was performed as follows: sex mismatches and individuals with abnormal numbers of sex chromosomes, related individuals, and those who withdrew consent. To avoid biases from population stratification, the sample was restricted to individuals of white British ancestry (as determined by self-report or analysis of genetic principal components (88)). Bringing together all the genetic and phenotypic data, including the cost data necessary to calculate IV models, resulted in 307,048 individuals included in the analysis. Further detail on these steps is provided in the Supplementary Material. Related individuals were analyzed separately for the within-family Mendelian Randomization analysis.

The most recent and largest genome-wide association study that did not explicitly overlap (91) with UK Biobank participants was Locke et al (92). Proxy SNPs were used for any SNPs identified in Locke et al (92) but not present in UK Biobank, provided that a suitable proxy with an R^2^ statistic between the proxy and missing SNPs of at least 0.8 was available in UK Biobank. To avoid violations of the IV assumptions due to linkage disequilibrium, only SNPs that were correlated with each other with an R^2^ of less than 0.001 within 10,000 kilobases were retained for analysis using the MR-Base R package (93).

In total, 79 of the 97 genome-wide significant SNPs identified in Locke et al (92) were included in the analysis, following this process and the removal of triallelic and unreconciled palindromic SNPs. Locke et al includes groups of heterogenous ancestry (94). The list of 79 SNPs from Locke et al included those from studies of both European and non-European ancestry. In sensitivity analysis, we re-ran the Mendelian Randomization analysis restricting the SNPs (n=69) from Locke et al to those of only European ancestry. Data on SNPs implicated in fat mass percentage used in multivariable analysis described below were taken from Lu et al (95).

Both the individual variants and a summary polygenic allele score created from these variants were used in analysis. The allele score was used in tests of association between potential omitted variables present at conception that were available in UK Biobank (sex, year of birth, month of birth) using linear regression. The allele score was calculated as the sum of the BMI-increasing alleles for SNPs attaining genome wide significance in Locke et al (92). Each SNP was weighted by the size of its effect on BMI.

We compared the Mendelian Randomization estimates to those from multivariable models by estimating the effect of a marginal unit of BMI on costs using a generalized linear model (GLM) with a gamma family and log link function following Dixon et al (86).^3^

The causal estimates from the Egger, median and mode estimators were converted from standard deviation units of BMI reported in the Locke et al (92) to natural units of BMI by dividing by the median standard deviation of BMI (4.6) in that study, as reported in Budu-Aggrey et al (96). This rescaling allows the results of all estimators to be interpreted as the marginal effect of a unit (kg/m^2^) increase in BMI on inpatient costs.

Analysis was conducted primarily in R using the MR Base package (93). Stata version 15.1 (StataCorp, College Station, Texas) was used for some elements of the analysis. Analysis code is available at github.com/pdixon-econ

## 4 Results

Of the 307,048 individuals included in the analysis sample, 54% were female (n=164,903), and mean age was 56.9 years (standard deviation: 8.0). Mean inpatient hospital cost per person-year of follow-up was £479, while median costs were £88. Mean and median follow-up of inpatient hospital data was 6.1 years. The most common ICD-10 chapters under which patients were admitted (other than for symptoms and findings not otherwise classified) were neoplasms (most commonly breast cancer) and musculoskeletal disorders (most commonly arthropathies).

There was evidence of association of the allele score with nine of the first ten principal components (largest p-value, from the eighth principal component=0.11)) but weaker evidence of association with month (p=0.464), year of birth (p=0.07) and sex (p=0.06). Sex and all ten principal components were included as covariates in all Mendelian Randomization models.

Results indicate that the effect of an additional unit of BMI is approximately 58% higher using IVW methods than under multivariable analyses (Table 1).

**Table 1.** Mendelian Randomization and multivariable estimates of marginal effect of an additional unit of BMI on per person year inpatient hospital costs.

|  | Beta (£) | SE | P Value |
| --- | --- | --- | --- |
| **Estimator** |  |  |  |
| Inverse variance weighted random effects estimator (IVW RE) | 21.22 | 3.50 | <0.001 |
| Multivariable estimate | 13.47 | 0.49 | <0.001 |

Note: We report p-values smaller than 0.001 as <0.001. Larger p-values are reported to two decimal places

However, there is evidence of heterogeneity (Cochran’s Q =107.8, p-value for null of no heterogeneity =0.01) in the base IVW results, one cause of which may be pleiotropy in violation of the exclusion restriction. Heterogeneity is apparent in the forest plot (Figure 1). A forest plot without heterogeneity would show all variants “lining up” around the same point estimate of effect, subject to sampling variation which will mean that not all variants would lie on precisely the same line even in the complete absence of heterogeneity.

**Figure 1.**
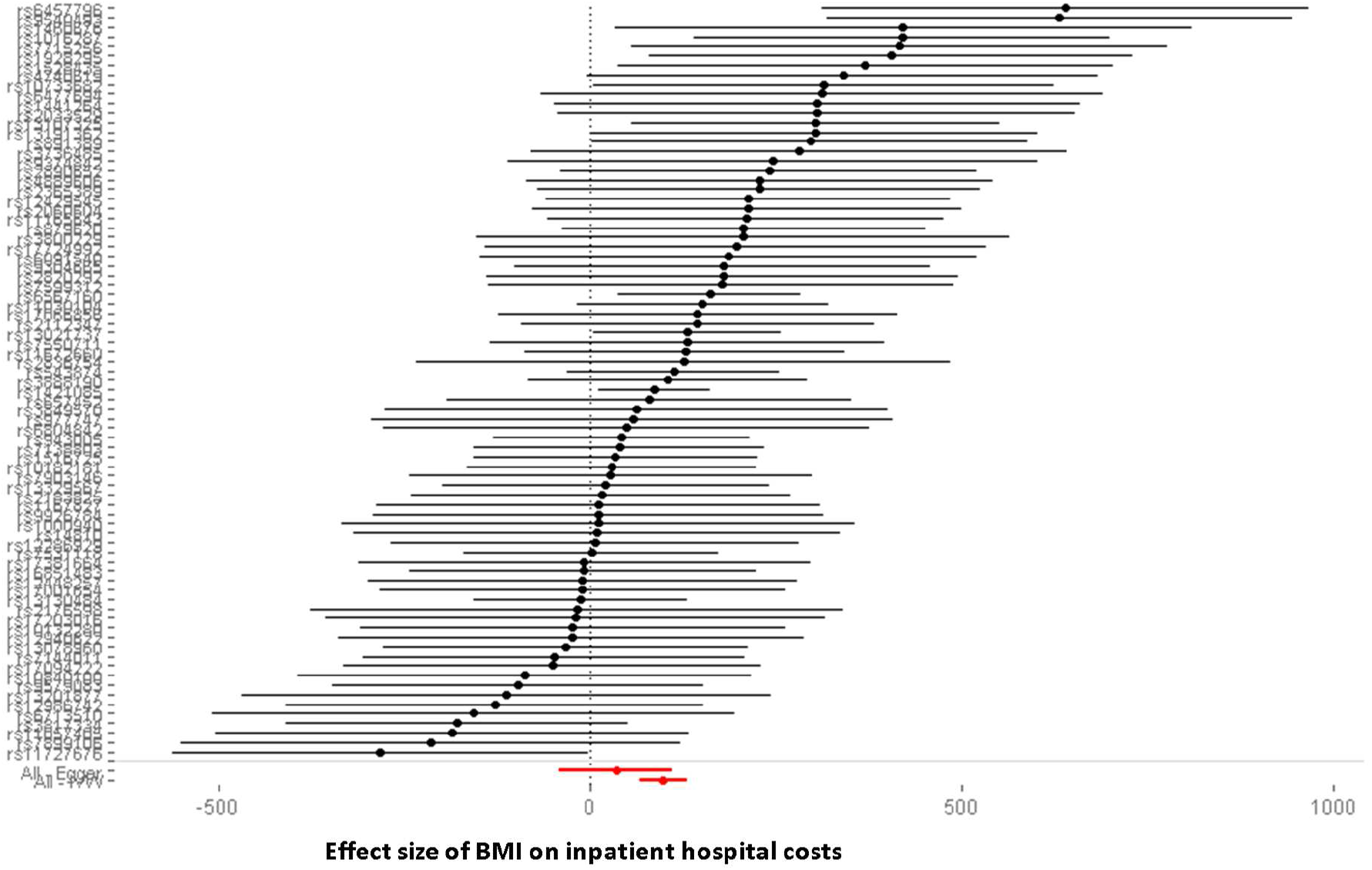
Forest plot of SNPs. The two diamonds at the bottom of the plot represent the point estimate the IVW estimate from using all variants together with a 95% confidence interval, and for contrast the MR-Egger point estimate with confidence intervals. The results of MR-Egger and other methods to adjust for pleiotropy are indicated in Table 2, again presented for comparison alongside the base IVW results.

**Table 2.** Results of primary Mendelian Randomization models.

|  | Beta (£) | SE | P-value |
| --- | --- | --- | --- |
| **Estimator** |  |  |  |
| IVW RE (for reference) | 21.21 | 3.50 | <0.001 |
| MR-Egger | 7.41 | 8.44 | 0.38 |
| Penalized weighted median | 18.85 | 5.00 | <0.001 |
| Weighted mode | 16.75 | 6.08 | 0.01 |

Figure 2 presents a scatter plot summarising these estimates. All estimators identify a positive effect of BMI on hospital costs, although the MR-Egger estimator is consistent with a null effect. The MR-Egger Cochran’s Q test of heterogeneity was 103.44 (p-value 0.02) and the intercept of this model was estimated as £1.93 (standard error: 1.07, p-value: 0.08). The IVW estimate is larger than all other estimates, although similar to the penalized weighted median estimate. If pleiotropy is present in the IVW model but not in the penalized weighted median model, it appears to be inflating somewhat the effect estimates, which would be the case if some of the included SNPs act on other conditions or traits that tend to increase inpatient costs on average.

**Figure 2.**
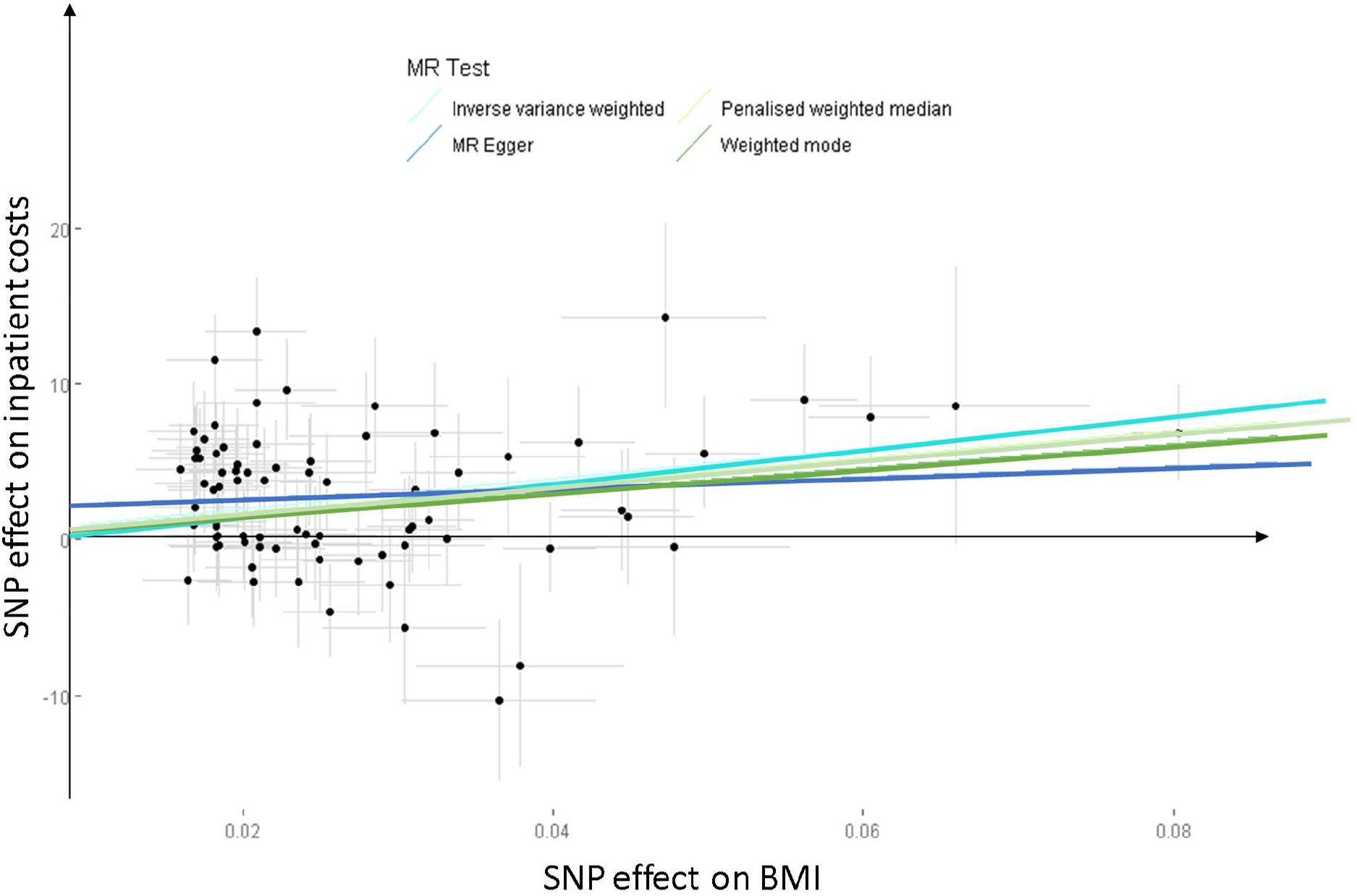
Scatter plot of estimators. The InSIDE assumption, which must be satisfied for unbiased MR-Egger estimates, is most likely to hold where the violations of the IV assumptions are caused by pleiotropy that does not influence omitted variables in the BMI-cost association. In practice, there is probably good reason to suspect violations of this type, as any variant that influences, for example, mental health may well be an omitted variable that independently influences both BMI and inpatient costs. In the case of this hypothetical example, instrument strength (measured by the association of BMI) may be correlated with a direct effect of the SNP (via mental health) on the cost outcome. Thus, any variant included amongst the 79 here that causes people to have inpatient care may well induce violations of InSIDE.

It is notable that the median and mode estimators are reasonably similar, despite the differences in the assumptions underlying each method. This is suggestive evidence that a similar causal effect is perhaps being identified by these two methods.

Models using a restricted set of SNPs indicated lower effects sizes and greater differences between the median and mode estimators (Table 3).

**Table 3.** Results of all Mendelian Randomization models with restricted SNP list.

|  | Beta (£) | SE | P-value |
| --- | --- | --- | --- |
| **Estimator** |  |  |  |
| IVW RE for reference | 18.70 | 3.80 | <0.001 |
| MR-Egger | 6.47 | 11.02 | 0.56 |
| Penalized weighted median | 16.10 | 5.03 | 0.001 |
| Weighted mode | 7.48 | 8.41 | 0.38 |

Heterogeneity was also somewhat lower when using the restricted list of SNPs, with Cochran’s Q for the IVW model of 88.32 (p-value=0.04).

### 4.1 Other sensitivity analyses

Using the full analysis sample, and dividing the IV-free BMI distribution into 100 quantiles, there was little evidence of non-linearity. There was evidence consistent with the null for a quadratic term (p=0.80), for differences in local average treatment effect estimates across quantiles (p=0.25)), for heterogeneity in the associations between the instrument and BMI across quantiles (p=0.09) and of a linear trend in the association between the instrument and BMI across quantiles (p=0.43). We conclude that the association between adiposity and inpatient hospital costs for this sample is approximately linear.

There was modest attenuation of the effect of BMI on costs when including body fat percentage in a multivariable Mendelian Randomization analysis. The causal coefficient on the body fat percentage IV was consistent with the null, while the effect estimate on BMI was within the confidence intervals of the base IVW estimate (Table 4).

**Table 4.** Results of multivariable Mendelian Randomization analysis.

|  | Beta (£) | SE | P-value |
| --- | --- | --- | --- |
| **Genetic variants** |  |  |  |
| IVW RE of BMI only (for reference) | 21.21 | 3.50 | <0.001 |
| BMI | 22.40 | 9.12 | 0.01 |
| Body fat percentage | -2.75 | 13.31 | 0.84 |

This suggests that any direct effect of body fat percentage on hospital costs is limited, and body fat percentage probably does not mediate the effects of body mass index on hospital costs. If body fat percentage were a mediator, the causal effect of BMI would change much more markedly between the univariable and multivariable Mendelian Randomization analyses.

Finally, application of the robust adjusted profile score method of Zhao et al (75) did not substantially alter the base IVW estimates of the causal effect.

**Table 5.** Results of robust adjusted profile score.

|  | Beta | SE | P-value |
| --- | --- | --- | --- |
| **Genetic variants** |  |  |  |
| Base IVW estimates (for reference) | 21.21 | 3.50 | <0.001 |
| Robust adjusted profile score | 21.69 | 3.06 | <0.001 |

Subject to the assumptions of the method, particularly that all pleiotropic effects have mean zero, this suggests that weak instruments and measurement error in the SNP-treatment association are not likely to be material sources of bias, at least for the base case results.

For the within-family analysis, 36,196 individuals were observed in 17,864 family units. The estimated effect of an additional unit of BMI was £3.46 (standard error 4.17, p-value 0.41, 95% confidence interval -£4.71 to £11.63). The effect size estimated is much smaller than in all other analyses and is consistent with the null. This could suggest that dynastic biases are responsible for the apparently material effect of BMI on inpatient costs in the main analysis using unrelated individuals. However, power to reject the null in this sample is weak, and it is possible that this finding is a false negative.

Finally, we consider disaggregation of all costs into elective costs, non-elective costs and other costs (Table 6). Effect sizes for an additional unit of BMI were larger in absolute terms for elective costs than for non-elective costs, and heterogeneity was more pronounced under the former (Cochran’s Q 107.2, p-value = 0.01) than the latter (Cochran’s Q, p-value = 0.12). These values were both lower than the corresponding value using all inpatient in the main analysis (Cochran’s Q=107.8, p-value=0.01), although the value of the statistic for inpatient-only costs was very similar to that for all costs. Heterogeneity for other costs only was similar to that for non-elective costs (Cochran’s Q=93.9, p-value=0.11).

**Table 6.** Disaggregated cost estimates.

|  | Beta | SE | P-value |
| --- | --- | --- | --- |
| **Cost outcome** |  |  |  |
| Base IVW estimates (all costs) | 21.21 | 3.50 | <0.001 |
| Elective costs only | 17.96 | 4.13 | <0.001 |
| Non-elective costs only | 9.00 | 3.80 | 0.02 |
| “Other” costs only | 4.17 | 3.13 | 0.18 |

The largest effect of BMI appears to be on elective care costs, for which estimated heterogeneity (as measured by Cochran’s Q) was similar to that for overall aggregate costs. While suggestive, there are a few reasons to be cautious in making this kind of interpretation. First, the categorisations used are somewhat arbitrary. Second, comparing the disaggregated costs both to each other and to all costs involves comparing different groups of individuals, since some cohort members report costs only in one subcategory of costs.

## 5 Discussion

The well-established positive association between adiposity and hospital costs appears to be causal. The results presented here using a novel Mendelian Randomization methodology suggest that this effect of a marginal unit of BMI is higher than that suggested by multivariable analyses. To our understanding, this is the first application of Mendelian Randomization to studying hospital costs as an outcome. Black et al ((28)) describe their biological relative instrumental variable analysis as Mendelian Randomization, but their work did not otherwise use data on genetic variants and is perhaps best considered as a distinct type of study design. Below, we briefly consider potential reasons for a narrower difference between IV estimates and multivariable estimates than is reported elsewhere, before considering the generalizability of the results, and finally potential remaining biases that may affect our estimates.

### 5.1 Comparison with other findings

Estimated differences between IV and multivariable models are smaller than those obtained from analyses using biological relatives as instruments, albeit these biological instrument studies were conducted on samples that may differ quite markedly from the sample studied here. Regression dilution (97), caused by measurement error in BMI, would tend to inflate differences between multivariable and IV analysis, since the multivariable results may be biased downward. One possible basis for the much narrower difference reported here is that the multivariable analyses are based on high-quality independent (i.e. not self-reported) measurements of weight and height.

There is a lack of a “gold standard” against which to judge multivariable and IV models or the various Mendelian Randomization estimators. Methods are being developed to choose amongst MR estimators including machine learning (55) and principled approaches to the treatment of “outlier” SNPs (98), although a degree of judgement and some contextual reasoning seems unavoidable in interpreting Mendelian Randomization analysis.

Despite the absence of a clear means to choose between types of estimator, policy evaluations and other quantitative analysis requiring estimates of the marginal cost of a unit of BMI should treat multivariable estimates as a lower bound. Analysts should consider including higher estimates of the cost of a marginal BMI unit in primary analysis.

### 5.2 Generalizability of findings

Are the results from this analysis likely to be generalizable to wider populations? Two issues merit consideration. The first is whether the Mendelian Randomization estimates are themselves helpful in understanding the effect of BMI on inpatient hospital costs. The second is whether the particular features of the UK Biobank sample, which is healthier and wealthier than the population from which it is drawn because of non-random participation, may itself create bias.

On the first point, Mendelian Randomization methods estimate, in this case, the effects on inpatient costs of a life-long exposure to BMI-increasing SNPs, rather than a temporary or acute effect of higher or lower BMI. We use the term “life-long” to refer to the effect of genetic variation determined at conception and assume that the association between the genetic variants and BMI does not change with age (99). The effect sizes estimated under all but the MR-Egger Mendelian Randomization analyses were larger in magnitude than the multivariable estimates, which suggests that they may reflect a cumulative exposure to higher BMI (100).

It is plausible that lifelong exposure to higher BMI, randomly determined at conception, could manifest in higher rates of inpatient admission and the use of more complex and expensive treatments amongst the middle-aged and early-old aged individuals represented in the UK Biobank cohort. As BMI is potentially modifiable, this suggests that policies targeting reductions in BMI (where clinically appropriate to do so) could reduce use of hospital resources (amongst other impacts on morbidity and mortality (101)).

The results of our analysis are most relevant to policy analyses effecting relatively modest changes in BMI, since the amount of variance in BMI explained by the SNPs used is less than 2%. We note that the more recent Yengo et al (91) GWAS explains a higher proportion of variance in BMI but has a substantial overlap with the UKBiobank, and therefore could not be used in this analysis. A further consideration is that included SNPs, and common genetic variants in general, may not satisfy the stable unit treatment value assumption in the sense that genetically influenced BMI may not have precisely the same impact on downstream cost outcomes than BMI influenced by drug therapies, diet interventions or exercise regimen (78). The difference in timing of effect between, for example, mid-life interventions targeting BMI and genetically elevated BMI (determined at conception) is another example of how Mendelian Randomization may not satisfy the stable unit treatment value assumption.

The second issue concerning the generalizability of our findings relates to the similarity or otherwise of the UK Biobank cohort to the wider population, and the implications that any differences may have on the generalisability of the results presented here. Relative to the UK population participants in the cohort study had lower levels of mortality (34), lower rates of health-compromising behaviour, and are better educated (81). BMI and use of hospital resources may themselves influence participation in the study (since sicker individuals were less likely to participate), and some degree of selection bias is possible (102). This specific bias goes by different names, including “collider bias” (103–105) and bias due to “bad controls” (106).

This selection appears to be problematic (in terms of bias and Type 1 error rates) for Mendelian Randomization only when selection effects are themselves particularly large (107). Since the size of this effect will generally be unknown (because the mechanism driving selection is unknown) it is not possible to be definitive about its scope in the present context. Gkatzionis and Burgess (108) suggest, on the basis of their simulations, that selection in general is probably less important as a source of bias than, for example, violations of the exclusion restriction caused by pleiotropy. It is also important to note that selection will also affect the non-causal multivariable estimates of a marginal unit of BMI presented alongside the causal IV analysis. It is possible that the precise figure for a marginal unit of BMI under either method may differ in other cohorts but nevertheless the ratio of the causal to non-causal costs will be stable when studied in other contexts.

### 5.3 Potential remaining biases

Three other potential sources of bias may be present. The first is bias from assortative mating, which refers to departures from random mating (109). The simulation and modelling study of Hartwig et al (110) found that bias from assortative mating would affect all forms of Mendelian Randomization analysis described above, including methods that attempt to account for pleiotropic SNPs.

Bias from assortative mating can overestimate SNPs-BMI and SNP-inpatient costs associations. This bias is larger when the strength of non-random assortment is high, the outcome is highly heritable and when the process of non-random mating has been present for a number of generations. In the absence of data relating to these influences, we simply note here that this bias may be present to some extent in the results presented here, and that data on family trios (parents and offspring) would help assess if assortative mating was present.

The second source of bias is from dynastic effects. We conducted a within-family Mendelian Randomization analysis to assess this bias. Results from these models were consistent with the null, but were much more imprecise than the other estimated models. The power offered by this analysis is a function of unknown variables, including the proportion of variance in offspring BMI that is explained by parental BMI. The dynastic bias is greater the larger is this effect. Evidence from Kong et al (79) find that the effect size of non-transmitted BMI-increasing alleles to be much smaller than the effect size for transmitted alleles (as modelled in the main analysis), which suggests dynastic effects may not be a large source of bias. Larger sample sizes, potentially involving meta-analysis across cohorts where within-family Mendelian Randomization is possible, would provide the one means to definitively understand whether power or substantive dynastic biases explain the null results from the within-family models estimated here.

Finally, Mendelian Randomization analysis may be confounded by cryptic geographic or population structure. There is some evidence, for example, that geographic structure is present in the UK Biobank sample (111). This could bias associations between health outcomes and genetic data, albeit that our inclusion of genetic principal components will address some but potentially not all such biases.

## 6 Conclusion

We have reported the first Mendelian Randomization analysis, using data on individual patients and specific SNPs, to estimate the causal effect of adiposity on inpatient hospital costs. Results suggest that multivariable analysis probably understates the effect of BMI on hospital costs. Mendelian Randomization is a feasible and potentially valuable form of analysis to answer these and related questions in health economics. The methods could be readily applied in modelling outcomes for other traits, behaviours, circumstances and diseases.

## Declarations

### Funding statement

PD, GDS, SH and NMD are members of the MRC Integrative Epidemiology Unit at the University of Bristol which is supported by the Medical Research Council and the University of Bristol (MC_UU_12013/1, MC_UU_12013/9). PD acknowledges support from a Medical Research Council Skills Development Fellowship (MR/P014259/1). SH was supported by Health Foundation grant “Social and economic consequences of health status - Causal inference methods and longitudinal, intergenerational data”

### Conflict of interest statement

The authors declare no conflicts of interest.

## Acknowledgments

The authors are grateful for comments on this work to seminar participants at Cambridge, Cornell, Newcastle and Oxford, and to conference participants at the Winter 2019 Health Economics Study Group meeting at York.

## Footnotes

1 Henceforth we use “multivariable” as shorthand for all study designs and estimators that do not use either formal randomization or reliance on some kind of natural experiment. We avoid the use of the term “observational” for this purpose, as Mendelian Randomization is itself a form of observational analysis. We also avoid the use of ordinary least squares (OLS) as shorthand for these other study designs, as many other

2 The treatment variable may be referred to as the exposure, the modifiable factor, the risk factpor or the intermediate phenotype. Following the econometrics literature, we will refer to BMI as the treatment variable.

3 The estimated effect of a marginal unit BMI differ between those reported in Dixon et al because the sample here is restricted to those with valid genetic data

