## Supplementary Material for "The causal effect of adiposity on hospital costs: Mendelian Randomization analysis of over 300,000 individuals from the UK Biobank"

March 26 2019

### 1 Introduction

This annex of Supplementary Material is arranged as follows:

- Section 2: Creation of elective, non-elective and other cost categories
- Section 3: Explanation of exclusions and participant numbers

#### 2 Creation of elective, non-elective and other cost categories

Total inpatient costs were disaggregated into elective, non-elective and other categories. These costs were calculated as follows.

First, elective, non-elective and other HRGs and their associated unit costs were identified from NHS Reference Costs. Second, these were linked to Finished Consultant Episode (FCE) output obtained from applying the NHS Grouper software to Hospital Episode Statistics data from the UK Biobank cohort. Where the link identified a FCE that associated with a HRG and unit cost that appeared in only one category, then this cost was assigned to that episode.

Second, elective care was identified if the admission method was coded as any of elective admissions from a waiting list, booked, planned, or coded as a transfer from another hospital other than in an emergency. Unit costs were then assigned according to this coding.

Some non-elective short stay care was coded as elective care. This was addressed by assigning costs on the basis of the share of elective/non-elective short stay care in total finished consultant bed days reported in NHS Reference Costs.

“Other” costs were calculated per individual by substracting elective and non-elective costs, as calculated above, from total costs.

### 3 Participant numbers

A total of 488,377 cohort participants were successfully genotyped. Analysis was restricted to those self-reporting “White British” ethnicity, or those who had very similar ancestral backgrounds, as determined by principal component analysis. This restricted the sample to 409,703 individuals. Individuals reporting sex mis-match and/or with sex chromosome aneuploidy were excluded (n=814). Kinship was estimated using the KING toolset (1), and identified 107,162 related pairs of individuals. An in-house algorithm removed individuals related to the greatest number of individuals until no related pairs remained, resulting in the exclusion of 79,448 individuals. Additionally, two individuals were related to a very large (more than 200) number of cohort members and were excluded. Finally, 76 individuals withdrew their consent for study participation and their details were removed. After these exclusions, 337,055 individuals remained in the dataset.

The in-house processing of the genetic data is described in more detail in Mitchell et al (2). Genetic data was also subject to  quality controls by UK Biobank (3).

The exclusions for the cost data proceeded as set out in Figure 1 of Dixon et al (4). Valid inpatient hospital cost data and body mass index data were available for 457,689 individuals. Bringing together the genetic and phenotypic (cost, BMI and other data) resulted in the final analysis group comprised of 307,048 individuals.
